# Redox-controlled structural reorganization and flavin strain within the ribonucleotide reductase R2b-NrdI complex monitored by serial femtosecond crystallography

**DOI:** 10.1101/2022.04.14.488295

**Authors:** Juliane John, Oskar Aurelius, Vivek Srinivas, In-Sik Kim, Asmit Bhowmick, Philipp S. Simon, Medhanjali Dasgupta, Cindy Pham, Sheraz Gul, Kyle D. Sutherlin, Pierre Aller, Agata Butryn, Allen M. Orville, Mun Hon Cheah, Shigeki Owada, Kensuke Tono, Franklin D. Fuller, Alexander Batyuk, Aaron S. Brewster, Nicholas K. Sauter, Vittal K. Yachandra, Junko Yano, Jan Kern, Hugo Lebrette, Martin Högbom

## Abstract

Redox reactions are central to biochemistry and are both controlled by and induce protein structural changes. Here we describe structural rearrangements and crosstalk within the *Bacillus cereus* ribonucleotide reductase R2b-NrdI complex, a di-metal carboxylate- flavoprotein system, as part of the mechanism generating the essential catalytic free radical of the enzyme. Femtosecond crystallography at an X-ray free-electron laser was utilized to obtain structures at room temperature in defined redox states without suffering photoreduction. We show that the flavin in the hydroquinone state is under steric strain in the R2b-NrdI protein complex, presumably tuning its redox potential to promote superoxide generation. Moreover, a binding site in close vicinity to the expected flavin O_2_-interacton site is observed to be controlled by the redox state of the flavin and linked to the channel proposed to funnel the produced superoxide species from NrdI to the di-manganese site in protein R2b. These specific features are coupled to further structural changes around the R2b- NrdI interaction surface. The mechanistic implications for the control of reactive oxygen species and radical generation in protein R2b are discussed.

## Introduction

Ribonucleotide reductases (RNRs) are essential metalloenzymes that employ sophisticated radical chemistry to reduce the 2’-OH group of ribonucleotides and thereby produce deoxyribonucleotides (dNTPs), the building blocks of DNA. This reaction is the only known pathway for the *de novo* synthesis of dNTPs. Three classes of RNRs are differentiated based on their structural features and radical generating mechanism. Class I consists of a radical generating subunit, R2 and a catalytic subunit, R1. Oxygen is required to generate a radical in the R2 subunit, which is reversibly shuttled to R1 to initiate the ribonucleotide reduction. Class I, found in eubacteria and all eukaryotes, is to date divided into five subclasses, Ia – Ie, based on the type of metal cofactor, metal ligands and radical storage in R2 (Högbom et al., 2020). The R2 subunit is characterized by a ferritin-like fold housing two metal ions coordinated by six conserved residues (Nordlund & Reichard, 2006), i.e. two histidines and four carboxylates (with the exception of subclass Ie (Blaesi et al., 2018; Srinivas et al., 2018)). The class Ib R2 subunit (R2b) metal site can bind two manganese ions or two iron ions. The two metal ions are oxidized from the M(II)/M(II) state to a short-lived M(III)/M(IV) intermediate which decays to M(III)/M(III) while producing a radical species on an adjacent tyrosyl residue (Tyr⋅), where it is also stored (Cotruvo et al., 2013; Cotruvo & Stubbe, 2010). The di-manganese R2b was shown to be the physiologically relevant form of R2b (Cotruvo & Stubbe, 2010; Cox et al., 2010); it accumulates higher amounts of radical and has higher enzymatic activity than the di-iron R2b. Molecular oxygen (O_2_) can directly activate the di-iron R2b (Huque et al., 2000). In contrast, the di-manganese R2b form cannot react with O_2_ but requires a superoxide radical (O_2_^•-^) provided by NrdI, a flavoprotein which is generally encoded in the same operon as R2b (Berggren et al., 2014; Cotruvo & Stubbe, 2010; Roca et al., 2008).

**Scheme 1.**
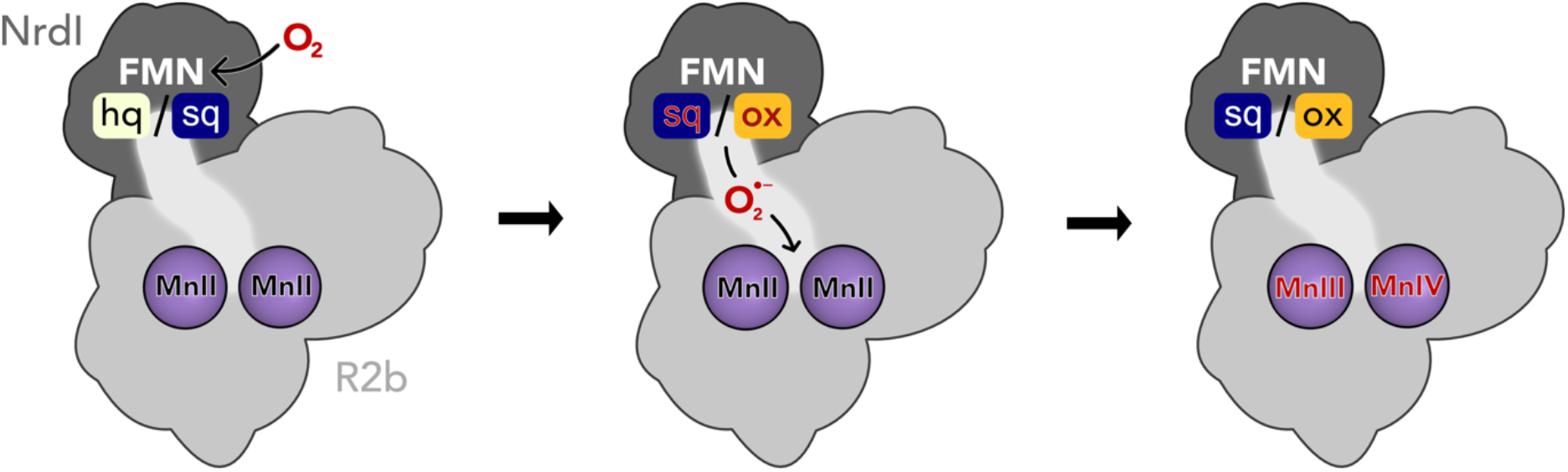
Activation of the di-manganese centre in ribonucleotide reductase class Ib R2. Both hydroquinone (hq) and semiquinone (sq) FMN of NrdI can reduce molecular oxygen to superoxide, which is shuttled to the metal site in R2b and activates the di-manganese cofactor. FMN is oxidized either to semiquinone or the fully oxidized form (ox) in the process.

NrdI is a small, globular protein that binds the redox-active flavin mononucleotide (FMN) cofactor. NrdI stands out in comparison to other flavodoxins, which perform single one- or two-electron transfers, by being able to perform two successive one-electron reductions (Cotruvo & Stubbe, 2008, 2010). Fully reduced NrdI with hydroquinone FMN (NrdI_hq_) can reduce O_2_ to O_2_^•-^ while being oxidized to semiquinone NrdI (NrdI_sq_) (Cotruvo et al., 2013). NrdI_sq_ can in turn produce a second O_2_^•-^ from another O_2_ molecule and become fully oxidized (NrdI_ox_) in the process (Berggren et al., 2014). The oxidation state of NrdI can be followed by ultraviolet–visible light absorption (UV–vis) spectroscopy. NrdI is faint yellow in the hydroquinone state, dark blue in the semiquinone state and bright orange in the oxidized state (Røhr et al., 2010) (Fig. S1). NrdI binds tightly to R2b in a 1:1 ratio with FMN at the protein–protein interface, forming a dimer of heterodimers. Previously obtained crystal structures of the R2b-NrdI complex from *Escherichia coli* and *Bacillus cereus* show that a conserved channel connects FMN and the metal site in R2b (Boal et al., 2010; Hammerstad et al., 2014). This channel is proposed to shuttle O_2_^•-^ produced by NrdI in complex with R2b to the metal site, where the radical is formed. Here we investigate the structural details and reaction mechanism of the activation of NrdI by O_2_ as well as the subsequent shuttling of O_2_^•-^ to the metal site for the generation of Tyr⋅ in R2b.

Studying structural changes in redox-active proteins is challenging from an experimental point of view. X-ray crystallography is a well-suited method for investigating both high-resolution structural details in the redox-active centres as well as overall reorganization of proteins. However, both metalloenzymes and flavodoxins are very sensitive to photoreduction during the exposure to the high-energy X-ray radiation of synchrotron sources. Synchrotron crystal structures of RNR and NrdI proteins invariably suffer from photoreduction, which complicates the determination of the oxidation state of redox active centres and has proven to be a problem for obtaining fully oxidized structures (Grāve et al., 2019; Johansson et al., 2010; Røhr et al., 2010). While there are well established experimental workarounds like short exposure times, helical data collection and serial synchrotron crystallography the time-scales necessary to obtain the diffraction data using these methods cannot fully avoid the reducing effects of the X-ray radiation (Spence, 2017). Serial femtosecond crystallography (SFX) overcomes this problem by the so-called diffraction-before-destruction principle (Doerr, 2011; Nass, 2019; Neutze et al., 2000). Short intense X-ray pulses of the duration of only a few femtoseconds are produced by an X-ray free electron laser (XFEL), illuminating one crystal and giving rise to one diffraction pattern at a time. The crystal is generally destroyed in the process and needs to be replaced by a new one for the next image. This method allows to record the scattering information before atomic displacement has time to occur resulting in a dataset effectively free from the effects of radiation damage (Spence, 2017).

Here we present the first SFX structures of R2b in complex with NrdI in the oxidized and hydroquinone state. The datasets reveal redox-dependent structural rearrangements both in the FMN binding pocket of NrdI and at the protein interface. Despite the significant rearrangement in the direct vicinity of the cofactor, FMN itself changes surprisingly little between redox states. This marks an interesting contrast to structures of free NrdI, which display significant conformational change of FMN between different oxidation states. We conclude that the R2b-NrdI complex formation is restricting FMN movement and inhibits this conformational change with implications for its redox potential and O_2_ reactivity. We also describe the first R2b-NrdI complex with a di-manganese metal centre from *Bacillus cereus*. The metal coordination in both structures blocks access to the channel that connects FMN and the metal site providing further information on gating and control of catalytic reactive oxygen species.

## Results

### SFX crystal structures of *Bacillus cereus* R2b in complex with oxidized and hydroquinone NrdI collected at ambient temperature

An initial crystallization condition for the R2b-NrdI complex was optimized to yield a sufficient amount of crystals smaller than 100 µm in the longest axis (Fig. S1), the maximum acceptable crystal size for the experimental setup. Crystals were initially tested at SACLA (SPring-8 Angstrom Compact free electron LAser, Japan) under aerobic conditions with a grease extruder setup (data not shown) (Sugahara et al., 2015; Tono et al., 2013). The crystals diffracted to 2 Å and proved to be stable under room temperature for several days and sturdy enough to handle the physical stress of being manipulated for the experiment. At LCLS (Linac Coherent Light Source at SLAC National Accelerator Laboratory, USA), we obtained two structures of the *Bacillus cereus* R2b-NrdI (*Bc*R2b-NrdI) complex in different defined redox states by SFX. The datasets were collected at room temperature under anaerobic conditions using the drop-on-demand sample delivery method (Fuller et al., 2017) (see Materials and Methods for details).

*Bc*R2b protein was produced metal-free to allow full control over metal loading during complex reconstitution. The metal content of the protein was determined by total-reflection X-ray fluorescence (TXRF) and only trace amounts of metals could be detected. The iron and manganese content per R2b monomer corresponded to metal-to-protein molar ratios of 0.27 ± 0.04%, and 0.07 ± 0.04%, respectively. The *Bc*R2b-NrdI complex was reconstituted *in vitro* by mixing both proteins in a molar ratio of 1:1 and set up for batch crystallization. Manganese was present both during the complex reconstitution and in the crystallization condition. Crystals for two different datasets were prepared to investigate different oxidation states of NrdI. The crystals for the first dataset were grown under aerobic conditions, yielding bright orange crystals, indicating that NrdI was fully oxidized (Fig. S1). The structure from these crystals is later referred to as *Bc*R2b_MnMn_-NrdI_ox_ (PDB ID: 7Z3D). The second dataset was obtained by reducing *Bc*R2b_MnMn_-NrdI_ox_ crystals chemically with sodium dithionite in an anaerobic environment. Consequently, the crystal colour changed from bright orange to faint yellow, indicating that NrdI underwent a two-electron reduction to the hydroquinone state (Fig. S1). The crystals were subsequently kept under anaerobic conditions, preventing reoxidation of NrdI. The corresponding structure is denoted *Bc*R2b_MnMn_-NrdI_hq_ (PDB ID: 7Z3E).

### Overall structure

The two structures were solved in space group *C*222_1_ and are of very similar quality with a resolution of 2.0 Å and similar unit cell dimensions (Table 1). The asymmetric unit contains one monomer of the 1:1 *Bc*R2b-NrdI complex. The physiological dimer can be generated by applying crystal symmetry (Fig. 1 A). Clear electron density maps allowed us to model residues 1-299 of 322 for R2b and 1-116 of 118 for NrdI in *Bc*R2b_MnMn_-NrdI_ox_ and 1-298 of R2b and all 118 residues of NrdI in *Bc*R2b_MnMn_-NrdI_hq._ Overall *Bc*R2b_MnMn_-NrdI_ox_ and *Bc*R2b_MnMn_-NrdI_hq_ are similar, as indicated by the Cα root-mean-square deviation (RMSD) value of 0.16 Å (Fig. 1 B). Refinement of both structures was conducted independently from each other following the same protocol (see Materials and Methods for details) with final *R*_*work*_/*R*_*free*_ of 0.16/0.20 for *Bc*R2b_MnMn_-NrdI_ox_ and 0.15/0.18 for *Bc*R2b_MnMn_-NrdI_hq_ (Table 1). Hammerstad *et al*. previously reported two crystal structures of the *Bc*R2b-NrdI complex (Hammerstad et al., 2014), obtained by single-crystal X-ray diffraction at a synchrotron source and under cryogenic conditions. Here, however, R2b harbours a di-iron active site. These structures will be referred to as *Bc*R2b_FeFe_-NrdI-1 and *Bc*R2b_FeFe_-NrdI-2 (PDB ID: 4BMO and 4BMP, respectively). Structural alignment between the published synchrotron and our XFEL structures shows that the overall fold of the complex is similar with Cα-RMSD values between 0.42 and 0.47 Å (Fig. 1 B). Compared to the *Bc*R2b_FeFe_-NrdI structures we observe a 7-8 residue extended ordered C-terminus in our structures. The C-terminus in *Bc*R2b_MnMn_-NrdI_ox_ continues in the same orientation as the C-terminal α-helix without keeping a helical conformation. It interacts with a groove between R2b and NrdI extending the R2b-NrdI binding area by forming hydrogen bonds to two other α-helices of R2b and a loop in NrdI (Fig. S2).

**Table 1.** Data collection and refinement statistics. Values in parenthesis are for the highest resolution shell.

|  | *BcR2b<sub>MnMn</sub>-NrdI<sub>ox</sub>* | *BcR2b<sub>MnMn</sub>-NrdI<sub>hq</sub>* |
| --- | --- | --- |
| PDB ID | 7Z3D | 7Z3E |
| **Data collection statistics** |  |  |
| XFEL source | LCLS MFX | LCLS MFX |
| Wavelength (Å) | 1.30 | 1.30 |
| Space group | *C*222<sub>1</sub> | *C*222<sub>1</sub> |
| Unit cell dimensions a, b, c (Å) | 61.4, 125.6, 145.0 | 61.2, 125.8, 144.8 |
| Unit cell angles α, β, γ (°) | 90, 90, 90 | 90, 90, 90 |
| Resolution range (Å) | 51.59 - 2.0<br>(2.034 - 2.0) | 51.47 - 2.0<br>(2.034 - 2.0) |
| Unique reflections | 38319 (1881) | 38188 (1880) |
| Multiplicity | 66.94 (25.42) | 59.92 (26.46) |
| Merged lattices | 14357 | 13509 |
| Completeness (%) | 99.91 (100) | 99.91 (100) |
| Mean I/sigma (I) | 3.247 (0.654) | 3.28 (0.791) |
| Wilson B-factor (Å<sup>2</sup>) | 35.45 | 33.22 |
| *R*<sub>split</sub> (%) | 11.1 (86.3) | 11.1 (76.5) |
| CC<sub>1/2</sub> | 0.989 (0.32) | 0.988 (0.39) |
| **Refinement statistics** |  |  |
| Resolution range<br>used in refinement (Å) | 24.17 - 2.0<br>(2.07 - 2.0) | 23.91 - 2.0<br>(2.07 - 2.0) |
| Reflections used in refinement | 38253 (3745) | 38119 (3729) |
| Reflections used for *R*<sub>free</sub> | 1909 (206) | 1897 (203) |
| *R*<sub>work</sub> (%) | 15.91 (30.40) | 15.16 (28.44) |
| *R*<sub>free</sub> (%) | 19.56 (32.27) | 18.37 (31.18) |
| RMSD, bond distances (Å) | 0.007 | 0.007 |
| RMSD, bond angles (°) | 0.79 | 0.81 |
| Ramachandran favored (%) | 99.27 | 98.79 |
| Ramachandran allowed (%) | 0.73 | 1.21 |
| Ramachandran outliers (%) | 0 | 0 |
| Rotamer outliers (%) | 0.79 | 1.05 |
| Clashscore | 1.86 | 2.58 |
| Protein residues (R2b + NrdI) | 415 (299 + 116) | 416 (298 + 118) |
| Average B-factor (Å<sup>2</sup>) | 47.27 | 43.66 |
| macromolecules | 47.24 | 43.64 |
| ligands | 34.66 | 30.38 |
| solvent | 50.57 | 47.09 |
| Number of non-H atoms | 3655 | 3652 |
| macromolecules | 3462 | 3477 |
| ligands | 33 | 33 |
| solvent | 160 | 142 |

**Figure 1.**
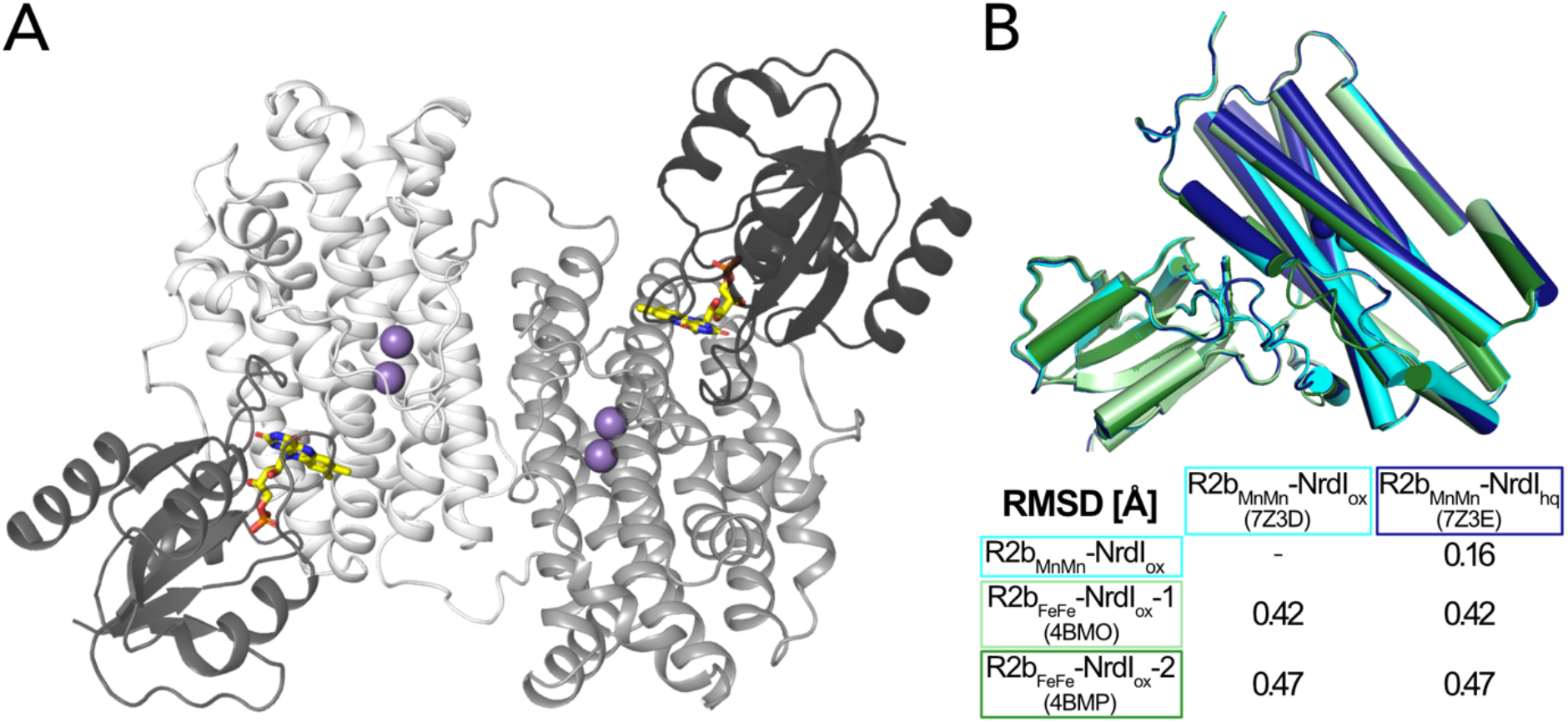
A) Structure of the *Bc*R2b-NrdI complex. (PDB ID: 7Z3D). In each 1:1 R2b-NrdI dimer, R2b and NrdI are coloured in lighter and darker grey, respectively. The second dimer is generated by crystal symmetry. Manganese ions are represented as purple spheres and FMN as yellow sticks at the R2b-NrdI interface. B) Superimposition of all structures of the *Bc*R2b-NrdI dimer with their corresponding Cα RMSD in the table. Individual structures are colour coded identically in the figure and the table.

### R2b complex formation prevents butterfly bend of FMN in NrdI

Three redox states of FMN are physiologically relevant for NrdI: oxidized (FMN_ox_), neutral semiquinone (FMN_sq_) and anionic hydroquinone (FMN_hq_) (Cotruvo et al., 2013; Røhr et al., 2010) (Scheme 2). Reduction of free FMN causes the isoalloxazine to bend along the N5-N10 axis leading to a “bufferfly bend” where the isoalloxazine moiety deviates from planarity (Fig. 2 A) (Zheng & Ornstein, 1996).

**Scheme 2.**
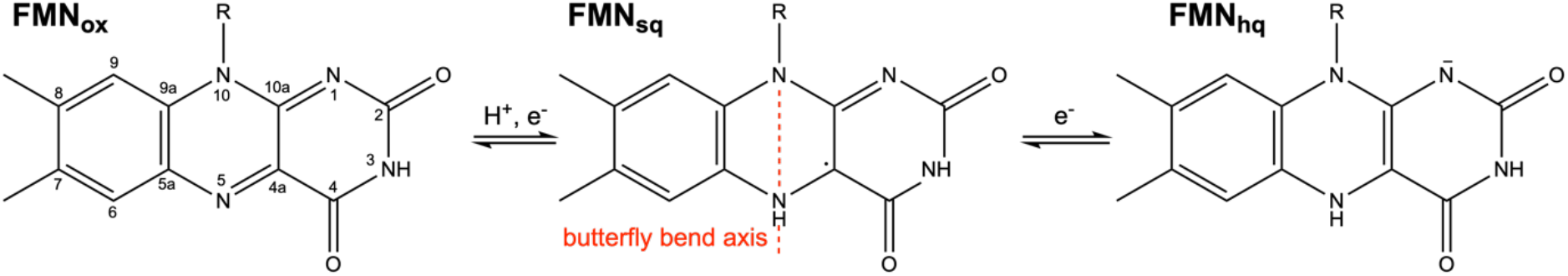
The physiologically relevant oxidation states of FMN in NrdI. The red dashed line marks the virtual axis between N5 and N10. The ribityl phosphate group is denoted as R.

**Figure 2.**
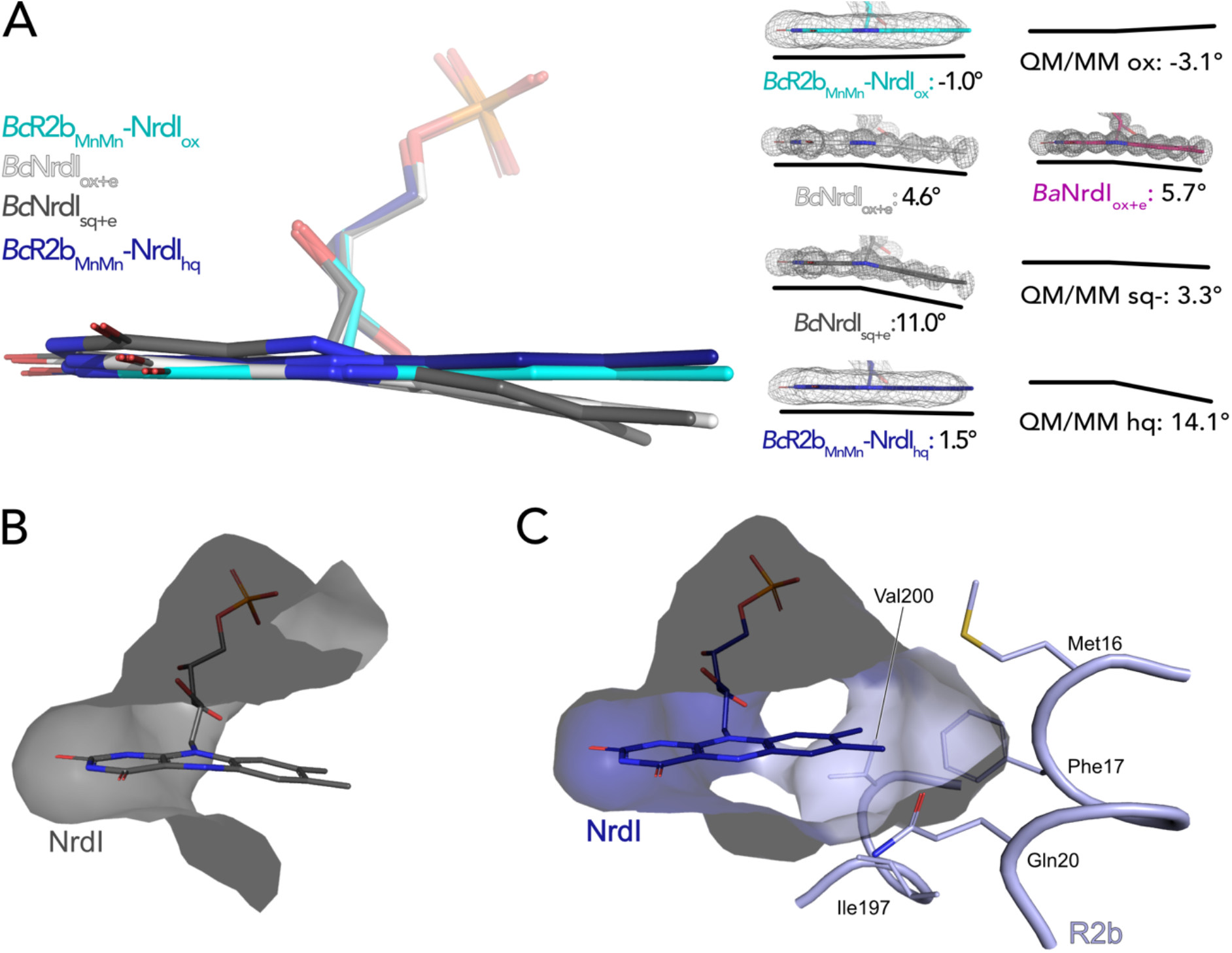
The butterfly bending conformations of FMN in different redox states. A) Left: Overlay of FMN cofactors of *B. cereus* NrdI in different crystal structures; Right: FMN cofactors in *Bc*NrdI and *Ba*NrdI with 2Fo-Fc maps contoured at 2 σ. Additionally, the theoretical calculated butterfly bend angles of FMN bound to *Bc*NrdI are shown (Røhr et al., 2010). Dark blue: *Bc*R2b_MnMn_-NrdI_hq_, cyan: *Bc*R2b_MnMn_-NrdI_ox_, grey: *Bc*NrdI_sq+e_ (PDB ID: 2X2P), white: *Bc*NrdI_ox+e_ (PDB ID: 2X2O), pink: *Ba*NrdI_ox+e_(PDB ID: 2XOD). B) The benzene ring of FMN in *Bc*NrdI_sq+e_ (PDB ID: 2X2P) is solvent exposed. C) In *Bc*R2b_MnMn_-NrdI_hq_ the FMN binding pocket is closed by residues contributed by R2b. Relevant residues are shown in sticks. The surface of the FMN binding pocket in NrdI in panels B) and C) is shown in grey.

NrdI binds one molecule of FMN non-covalently. The pyrimidine ring of the 7,8-dimethyl-isoalloxazine ring is buried and tightly bound to the protein by a combination of hydrogen bonds and π-stacking interactions while the benzene ring and part of the phosphate tail are solvent exposed (Fig. 2 B). The crystal structure of *Bc*NrdI has been described previously by Røhr et. al and shows the FMN-binding pocket on the protein surface (Røhr et al., 2010). The same study also presents quantum mechanics/molecular mechanics (QM/MM) calculations of the theoretical butterfly bend for FMN bound to NrdI. The calculated angles are shallow with -3.1° for FMN_ox_ and -3.9° for FMN_sq_ and more pronounced with 14.1° for FMN_hq_ (Fig. 2 A). The authors investigated the influence of photoreduction on the butterfly bend in FMN bound to *Bc*NrdI. The crystals for the first structure (PDB ID: 2X2O, 1.1 Å resolution) were produced from oxidized protein. The measured butterfly bend in this structure is 4.6° and does thus not correspond well to the calculated FMN_ox_ or FMN_sq_ angle. It does however resemble the calculated theoretical butterfly bend of 3.3° of the physiologically not relevant anionic semiquinone state of FMN (FMN_sq-_) (Fig 2 A). The second NrdI structure (PDB ID: 2X2P, 1.2 Å resolution) structure was produced from protein in the semiquinone state and the butterfly bend was 11° after data collection, similar to the calculated FMN_hq_ angle (Fig. 2 A). The authors could confirm the reducing effect of the synchrotron radiation on FMN by comparing Raman spectra of the crystals before and after data collection. These structures will be referred to as *Bc*NrdI_ox+e_ and *Bc*NrdI_sq+e_ to emphasize the photoreduction. Johansson et al. observed the same discrepancy between the butterfly bend for a synchrotron structure of initially oxidized NrdI from *Bacillus anthracis* (*Ba*NrdI), a protein with 99% sequence identity to *Bc*NrdI and identical FMN protein environment (Johansson et al., 2010). *Ba*NrdI_ox+e_ (PDB ID: 2XOD, 1.0 Å resolution) is even more bent than *Bc*NrdI_ox+e_ with 5.7°, indicating significant photoreduction of FMN during data collection. Figure 2 A lists the FMN angles of the calculated and measured structures.

The use of SFX allowed us, for the first time, to investigate the conformation of oxidized FMN in NrdI. In *Bc*R2b_MnMn_-NrdI_ox_, the isoalloxazine moiety of FMN_ox_ is almost planar, with a butterfly bend of -1°. This value is comparable to the angle of -3.1° for FMN_ox_ in NrdI calculated by QM/MM (Røhr et al., 2010). Unexpectedly, the bend of FMN_hq_ in the *Bc*R2b_MnMn_- NrdI_hq_ structure is minimal with only 1.5 ° and thus similar to FMN_ox_ in the *Bc*R2b_MnMn_-NrdI_ox_ structure (Fig. 2 A). Additionally, we calculated an isomorphous difference (Fo(ox)-Fo(hq)) map of *Bc*R2b_MnMn_-NrdI_ox_ and *Bc*R2b_MnMn_-NrdI_hq_ using the phases of the oxidized dataset. These maps reduce model bias by directly comparing the experimental data and are very sensitive to subtle changes of atom positions in different data sets (Rould & Carter, 2003). The Fo(ox)-Fo(hq) map shows a slight movement of the oxygen on C4 of FMN indicating a small twist of the pyrimidine ring between both structuresbut no further bending (Fig. 3). This angle of *Bc*R2b_MnMn_-NrdI_hq_ does not correspond to either the calculated bend of FMN of 14.1° nor the experimental bending angle of *Bc*NrdI_sq+e_ of 11°. In the R2b-NrdI complex the solvent exposed part of FMN is covered by R2b, which closes the binding pocket. The residues Met16 Phe17, Ile197 and Val200 of the R2b subunit interact hydrophobically with the benzene ring, while Gln20 is sterically hindering it from bending (Fig. 2 C). Together they form a rigid binding pocket for the isoalloxazine moiety of the cofactor, thereby preventing FMN_hq_ from bending.

**Figure 3.**
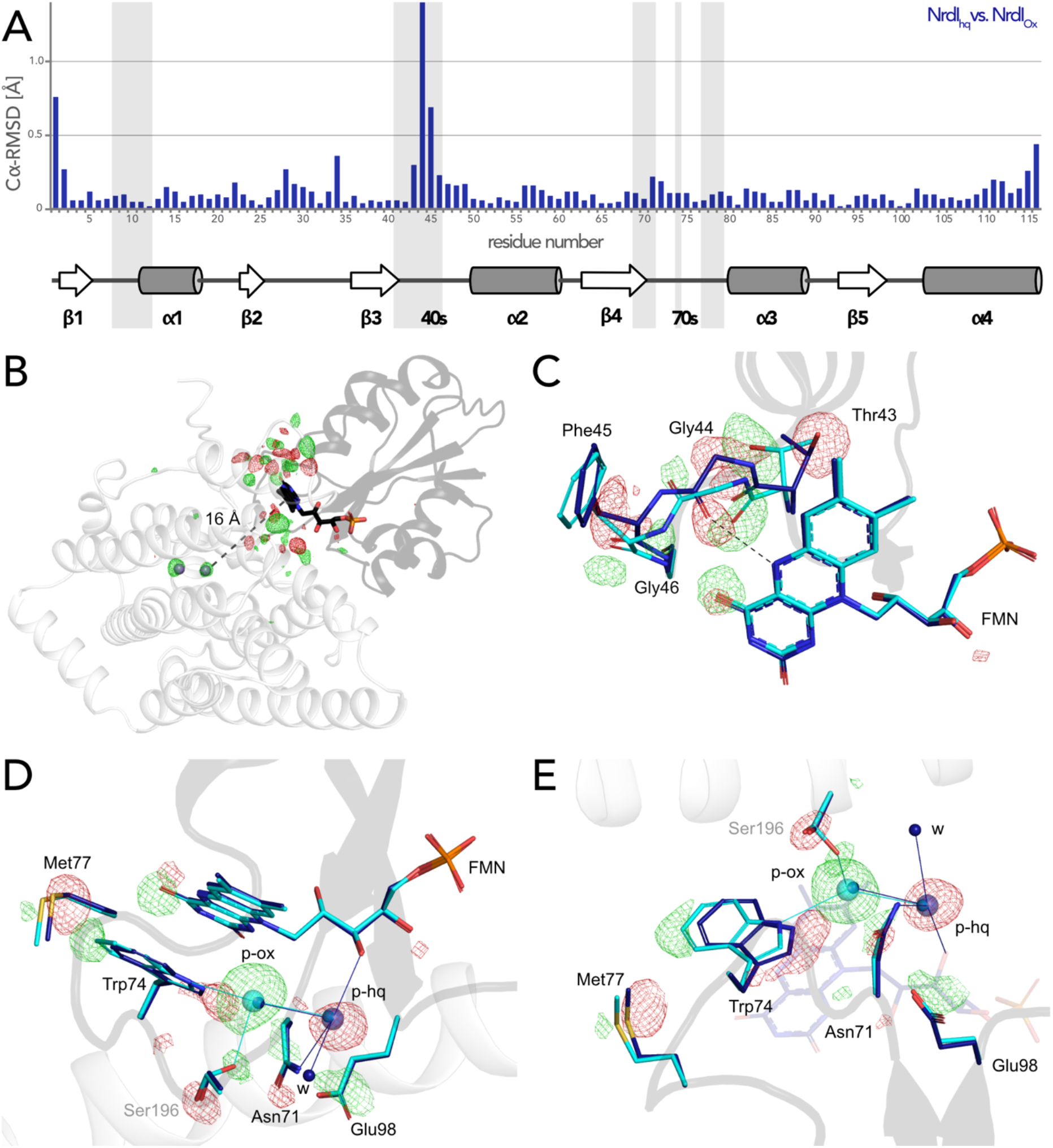
FMN environment in the oxidized and hydroquinone *Bc*R2b-NrdI. A) Structural alignment of NrdI in *Bc*R2b_MnMn_-NrdI_ox_ compared to *Bc*R2b_MnMn_-NrdI_hq_. Cα-RMSD in Å are shown for each residue of NrdI between the two models. The secondary structure assignment corresponding to the residues of NrdI is represented in the cartoon below. NrdI interactions with FMN are marked with light grey background B) Fo(ox)-Fo(hq) map contoured at 4.5 σ for the *Bc*R2b-NrdI complex with positive density in green and negative density in red. FMN are shown in sticks, manganese ions as purple spheres, R2b as transparent cartoon in white and NrdI in transparent black. Differences between both datasets cluster around the FMN at the R2b/NrdI interface. The distance between metal site and FMN is around 16 Å, marked with a dashed line. C) + D) + E) Close-ups of the difference density around the FMN from different angles including superposition of *Bc*R2b_MnMn_-NrdI_hq_ in dark blue and *Bc*R2b_MnMn_-NrdI_ox_ in cyan. Fo(ox)-Fo(hq) map contoured at 4.5 σ for all panels. Adjacent secondary structure elements are shown as transparent cartoon in white for R2b and black for NrdI. C) Rearrangement of the 40s loop of NrdI. Gly44 flips 180° in the hydroquinone state and forms a hydrogen bond with N5 of FMN (shown as dashed line). D) Rearrangements of side chains on the R2b facing side of FMN. Residues of NrdI are labelled in black, Ser196 from R2b in light grey. An unexplained density, larger than water, moves between the structures, named p-ox and p-hq and is marked as transparent big spheres; waters are represented as small opaque spheres. E) Same side chains as in D) are shown at a different angle facing the FMN. FMN is shown transparent in the background for clarity.

### Reorganization of FMN environment and binding position controlled by FMN redox state

The change of redox state of FMN causes movement around the cofactor in both proteins. Alignment of the NrdI backbone of the *Bc*R2b_MnMn_-NrdI_ox_ and the *Bc*R2b_MnMn_-NrdI_hq_ structures shows a clear reorganization in the 40s loop close to the isoalloxazine ring of FMN (Fig. 3 A). The biggest change can be seen for Cα of Gly44 and a smaller shift for Cα of Thr43 and Phe45. The reorganization of the 40s loop of NrdI after reduction is also apparent in the superposition of both structures and in the Fo(ox)-Fo(hq) map (Fig. 3 B and C). Notably, Gly44 of NrdI is flipped by 180° and forms a hydrogen bond with the hydrogen of N5 on FMN. The hydrogen bond between O4 of FMN and Gly46 shifts slightly. Thr43 in turn is shifted towards FMN and its side chain is rotated by 180°; Phe45 is also slightly rotated (Fig. 3 C). This redox-dependent conformational change of residues 43 to 45 was previously also observed in crystal structures of *Bc*NrdI and *Ba*NrdI in absence of the R2b subunit (PDB ID: 2X2O, 2X2P, 2XOD, 2XOE) (Johansson et al., 2010; Røhr et al., 2010).

Redox-induced changes are also visible on the side of the isoalloxazine ring facing R2b. An unknown molecule is bound within 5 Å from the reactive C4a of FMN in *Bc*R2b_MnMn_-NrdI_ox_, which binds O_2_ to reduce it to O_2_^•-^ (Ghisla & Massey, 1989). The molecule is 3- coordinated to Trp74 (NrdI), Ser196 (R2b) and a water (p-ox in Fig. 3 D and E). A similar density was previously observed in the di-iron *Bc*R2b-NrdI synchrotron structures where it was modelled as a chloride ion. In our structure, placing a water in this position does not sufficiently explain the density while modelling it as a chloride ion leaves no residual density (Fig. S3). Chloride is abundant in the crystallization condition; however, serine and tryptophane are not typical chloride ligands (Carugo, 2014). The exact nature of the molecule could not be determined and was thus left as a water in the final model. Interestingly, the binding position undergoes a redox dependent switch: In the reduced structure the density is instead found at a position between the oxygen on C2 of FMN_hq_ and Asn71 of NrdI, 7 Å from the reactive C4a. Two adjacent water molecules make the unknown molecule 4-coordinated (p-hq in Fig. 3 D and E). To accommodate for the change of position residues Met77, Trp74, Asn71 and Glu98 from NrdI and Ser196 from R2b move in a plane parallel to the FMN isoalloxazine ring (Fig. 3 D and E). The unidentified molecule exchanges position with two different water molecules of the well-ordered solvent network connecting FMN with the active site in R2b. The solvent network is housed by a channel between both proteins lined by the side chains of Ser162, Tyr166, Lys263 and Asn267 and the mainchain of Glu195 and Ser196 (Hammerstad et al., 2014). The position of the density in *Bc*R2b_MnMn_-NrdI_ox_ is close to the channel entrance on the R2b surface marked by Lys263, Asn267 and Ser196 (Fig. S4). Despite the large structural rearrangements around the FMN, the Fo(ox)-Fo(hq) map clearly shows that no redox-induced movement protrudes further down the channel and that the coordination of the metal site is unaffected by the change of NrdI redox state (Fig. 3 B and S4).

### Access to the metal site is not gated by FMN oxidation state, complex formation or manganese binding

Both SFX *Bc*R2b-NrdI structures display a di-manganese metal centre with similar coordination. In *Bc*R2b_MnMn_-NrdI_ox_ the two metal ions of R2b are refined at full occupancy and the coordination sphere is clearly defined in the electron density map (Fig. 4 A). Mn1 is 4- coordinated by His96, Glu93, Asp62 and Glu195. Mn2 is 5-coordinated by His198, Glu93, Glu195 and Glu161. Notably, Glu195 and Glu161 exhibit two alternative conformations each and alternately coordinate Mn2 in a monodentate or bidentate fashion. Importantly, while Glu195 bridges the two ions, Glu161 coordinates only Mn2. No coordinating waters could be identified (Fig. 4 B). The protein complex used in the crystallization for the *Bc*R2b_MnMn_-NrdI_ox_ dataset has not undergone any oxidation of the metal site and both manganese ions are in the Mn(II)/Mn(II) oxidation state (see Materials and Methods and Discussion for details). Notably, the reduction of FMN within NrdI occurs at about 16 Å from the metal site and does not affect the metal coordination in *Bc*R2b-NrdI (Fig. 3 A). The Fo(ox)-Fo(hq) map shows slightly lower occupancy for both manganese ions in *Bc*R2b_MnMn_-NrdI_hq_ (Fig S5). The loss of metals can be explained by the reduction treatment of the crystals for this dataset and Mn1 was modelled with 80%, Mn2 with 90% occupancy in the reduced structure. Both structures show the same metal-metal distance of 3.8-3.9 Å within experimental error for this resolution (Fig. 4 B).

**Figure 4.**
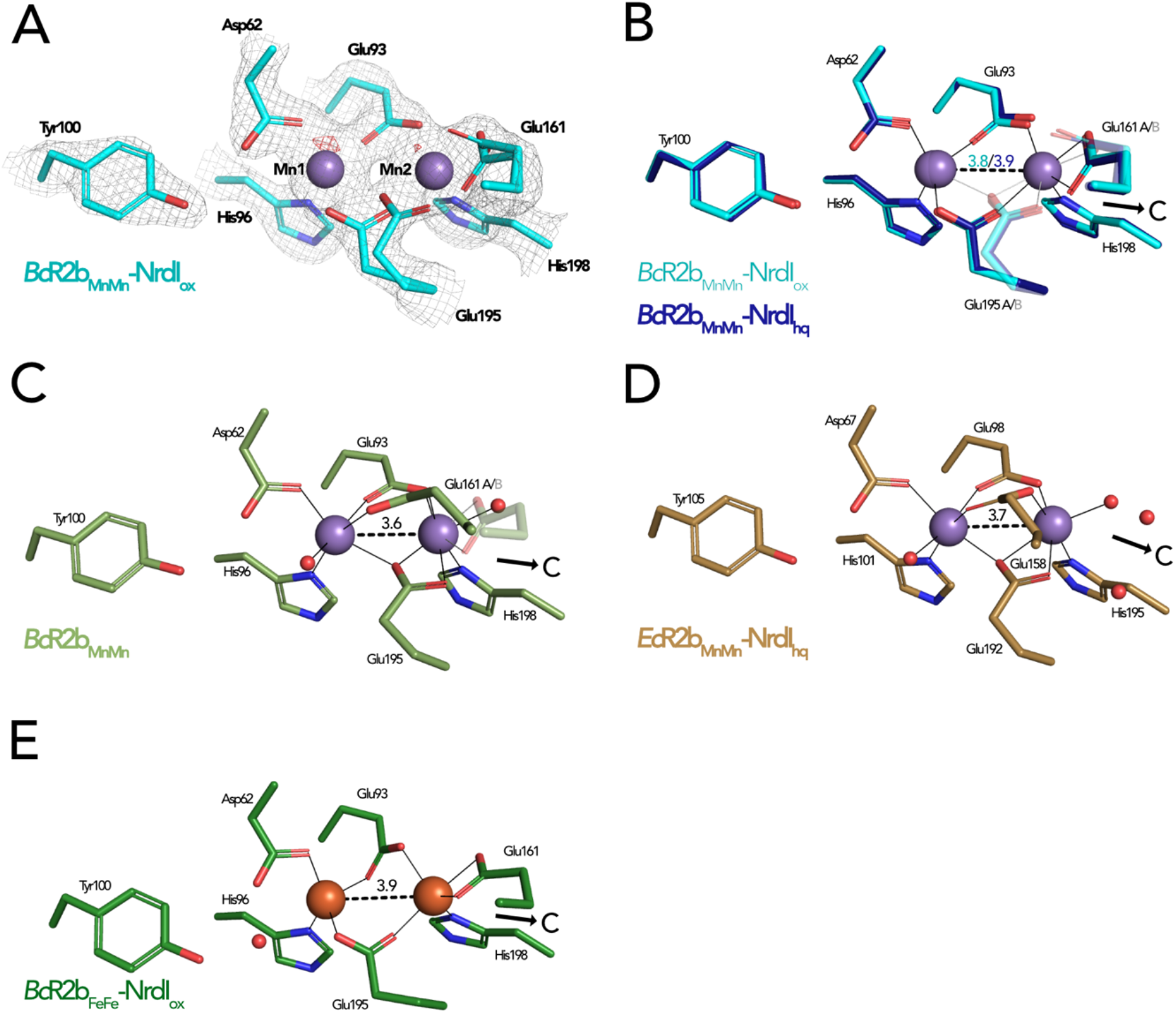
Comparison of active site architecture in different R2b structures including the radical harbouring tyrosine. Manganese ions are shown as purple, iron ions as orange and waters as red spheres. The direction of the connecting channel between R2b and NrdI is indicated by a black arrow and “C”. A) *Bc*R2b_MnMn_-NrdI_ox_ active site. The 2Fo-Fc map is contoured at 2 σ shown as grey mesh and Fo-Fc is contoured at 4 σ with negative density in red (no positive density is present). B) Superposition of *Bc*R2b_MnMn_-NrdI_hq_ and *Bc*R2b_MnMn_-NrdI_ox_. The metal coordination is identical for both structures. C) Active site of *Bc*R2b_MnMn_ (PDB ID: 4BMU). The metal site of chain A is shown with Glu161 in both the open (opaque) and closed (transparent) conformation. D) Active site of *Ec*R2b_MnMn_-NrdI_hq_ (PDB ID: 3N3A). Glu158 is in the open conformation and two waters are in the position of the Glu161 of *Bc*R2b. E) Active site of the di-iron *Bc*R2b-NrdI complex (PDB ID: 4BMP). The metal coordination is similar to the manganese containing structures with Glu161 in the closed conformation. An additional water is hydrogen-bonded to Asp62 and Tyr100. Metal-metal distances are shown as dashed and metal ligands as solid lines; residues in alternative conformations are shown transparent for panels B)-E).

The metal-ligand Glu161 is likely a key player in the activation of the di-manganese centre as it is proposed to gate the access to the metal site for the oxidant produced by NrdI (Boal et al., 2010). Two main conformations have been observed for this ligand: In the “closed” conformation the glutamate interacts only with Mn2 and is proposed to prevent O_2_^•-^ from reaching the metal site by obstructing the channel. In the “open” conformation, on the other hand, Glu161 bridges both manganese ions leaving space for three water molecules to connect Mn2 to the water network in the channel between FMN and the metal site. The open conformation of the equivalent glutamate has been observed in structures of di-manganese R2b alone, e.g. from *B. cereus, E. coli* and *Streptococcus sanguinis* (PDB ID: 4BMU, 3N37, 4N83) (Fig. 4 C) but also in the R2b-NrdI complex of *E. coli* (PDB ID: 3N3A) (Fig. 4 D) (Boal et al., 2010; Hammerstad et al., 2014; Makhlynets et al., 2014). Thus, NrdI binding to R2b is not responsible for triggering the open conformation of Glu161 (Glu158 in *E. coli* and Glu157 in *S. sanguinis*). It has been hypothesized that the presence of manganese ions in the active site could cause the glutamate to shift to the open conformation as the closed conformation is typically observed in R2b structures containing a di-iron site, e.g. in the *Bc*R2b_FeFe_-NrdI complex (PDB ID: 4BMP) (Fig. 4 E) (Hammerstad et al., 2014). Indeed, since the di-iron form of R2b is oxidized by O_2_ via a NrdI-independent pathway, the movement of the glutamate is not required for radical generation. However, our SFX *Bc*R2b_MnMn_-NrdI structures harbour a di-manganese site and exhibit a Glu161 in the closed conformation, thereby demonstrating that manganese in the active site is not sufficient to induce the glutamate shift. Finally, our radiation damage free structures also show that the opening of the channel towards the metal site is independent from the FMN oxidation state, as Glu161 is in the closed conformation when di-manganese R2b is in complex with either NrdI_ox_ or NrdI_hq_ (Fig. 4 B). Altogether, our data shows that the glutamate shift of Glu161 is not caused by the presence of manganese ions, R2b-NrdI complex formation, a specific NrdI redox state or a combination of these factors.

## Discussion

In this study we present two structures of the manganese containing R2b-NrdI complex of *Bacillus cereus* with NrdI in two different oxidation states. These structures were obtained by room temperature SFX data collection using XFEL radiation in contrast to all previously studied R2b or NrdI structures, which were obtained by classical single-crystal synchrotron data collection at cryogenic temperatures. Determination of the oxidation state of redox active enzymes via synchrotron radiation is problematic because exposure to X-rays exerts photoreduction on redox centres. Investigation of artifact-free oxidized enzymes is therefore exceedingly challenging with synchrotron radiation. In the context of class Ib RNRs this affects both the redox active metal centre of R2b and FMN cofactor of NrdI. SFX eliminates the effects of photoreduction observed during synchrotron-based data collection. We describe SFX structures of the R2b-NrdI complex with NrdI in the oxidized and hydroquinone oxidation state. Both 2.0 Å structures are of similar quality as the previously published synchrotron structures of the same complex with a di-iron centre (Hammerstad et al., 2014) showing that the change of methodology does not affect the quality. This allowed detailed examination of structural reorganization induced by changes in FMN redox state. Notably, the FMN conformation *per se* changes little between the oxidized and hydroquinone oxidation state in R2b-NrdI. This is surprising since it was previously shown that FMN is bent in reduced NrdI when not bound to R2b. The complexation of R2b and NrdI thus prevents FMN bending and exerts strain on the isoalloxazine ring in the hydroquinone state. Other flavoproteins have been shown to tune the redox potential of FMN by forcing it into a specific binding angle (Senda et al., 2009; Walsh & Miller, 2003). A recent study by Sorigué et al. presented a SFX structure of the flavoenzyme fatty acid photodecarboxylase (FAP) in complex with an oxidized flavin cofactor exhibiting a butterfly bending angle of 14° (Sorigué et al., 2021). Using time resolved SFX and complementary approaches, the study showed that during the enzymatic cycle of FAP the conformation of the flavin retains the butterfly bend in the different redox states. The authors conclude that the butterfly bend is enforced by the protein scaffold to promote flavin reduction. Following the same line of reasoning we propose that R2b binding to NrdI restricts FMN bending and thus, opposite to FAP, changes the FMN redox potential of the hydroquinone to favour FMN oxidation and reduction of molecular oxygen to superoxide (Scheme 3). This notion is supported by a study by Cotruvo et al. demonstrating that superoxide production by the *in vitro* R2b-NrdI_hq_ complex of *Bacillus subtilis* is about 40 times faster than the production of superoxide by free NrdI_hq_ under the same experimental conditions (Cotruvo et al., 2013). Controlling the superoxide production by complex formation could serve as a mechanism to protect the cell from production of superoxide by free NrdI. The binding of R2b may also facilitate access of O_2_ to FMN by forming a hydrophobic binding pocket around the benzene ring of FMN. O_2_ is reduced to superoxide at the C4a atom of the isoalloxazine ring of FMN and we observe a molecule bigger than a water bound about 5 Å away from the C4a atom in the oxidized R2b-NrdI complex. From the experimental setup it seems unlikely that the unidentified molecule is a superoxide ion because FMN never underwent a redox-cycle in this structure. However, refinement with a superoxide ion leaves little residual density, so the binding could fit a molecule of its size (Fig. S3). The density could thus represent a potential binding position of superoxide after its generation (Fig. S3). In addition, the density undergoes a redox-dependent switch of position moving further away from the reactive C4a (7 Å) towards the channel entrance in the reduced R2b-NrdI complex. This move is accompanied by a change of the putative coordination sphere. Importantly, in both oxidized and reduced states, the unknown molecule is integrated into the conserved hydrogen-bonding network connecting FMN and the metal site and switches position with a water. Even though the exact nature of the observed electron density is unknown we conclude that the binding properties of these key positions are controlled by the FMN redox state and could mark the route superoxide would travel after being produced by the FMN.

**Scheme 3.**
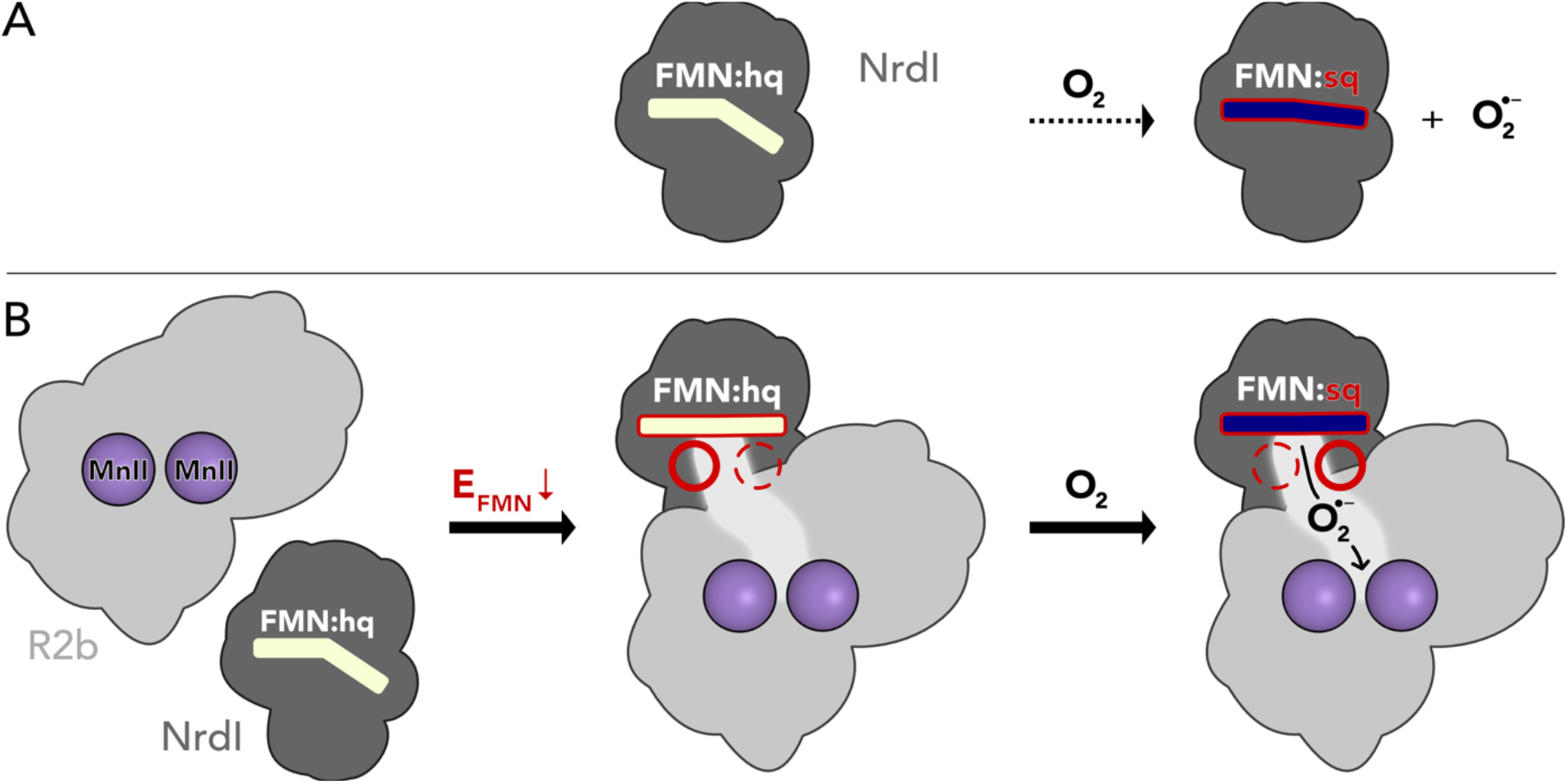
Proposed mechanism for the assembly of the R2b-NrdI complex. A) Free NrdI_hq_ and molecular oxygen react slowly to produce superoxide and NrdI_sq_. B) R2b and NrdI form a complex which imposes strain on the FMN bend and lowers its redox potential to favour oxygen reduction. The complex formation also generates a redox state-controlled potential superoxide binding site close to the reactive carbon of FMN. Upon exposure to molecular oxygen, superoxide is generated in the complex and shuttled towards R2b.

Taken together, our results suggest that the production of superoxide by NrdI and the radical generation in R2b is elegantly orchestrated by the formation of the R2b-NrdI complex as well as redox state control of binding positions (Scheme 3). R2b-NrdI binding induces conformational strain in the flavin, lowering its redox potential and thus promoting superoxide generation. The interaction surface also provides binding positions that are controlled by the redox state of the flavin, presumably involved in gating of channel and metal site access.

## Materials and Methods

### Protein Expression

The plasmids containing the genes for both *Bc*R2b (pET22b-Bc*r2b*) and *Bc*NrdI (pET22b- Bc*nrdI*) were kindly provided by Marta Hammerstad and Kristoffer Andersson (University of Oslo). The protein expression of *Bc*NrdI was adapted from Røhr et al. (Røhr et al., 2010). *E. coli* BL21(DE3) (New England Biolabs, Frankfurt am Main, Germany) cells were transformed with the pET22b-Bc*nrdI* plasmid. A preculture of Lysogeny Broth medium (Formedium, Norfolk, UK) was inoculated with a single colony, grown over night at 37 °C and 200 rpm. The next day large-scale cultures of 1.6 L Terrific Broth medium (Formedium) per glass bottle supplemented with 100 µg/ml carbenicillin (Alfa Aesar, Kandel, Germany) and 1:10000 v/v antifoam 204 (Merck, Darmstadt, Germany) were inoculated with 0.5% v/v of the preculture. The cells were incubated in a LEX bioreactor (Epiphyte3, Toronto, Canada) at 37 °C until an optical density at 600 nm of about 0.8 was reached. The cultures were cooled down to 20 °C, and the protein expression was induced with 0.8 mM isopropyl β-D-1-thiogalactopyranoside (IPTG) (Formedium). The cells, harvested after 12-16 hours of expression by centrifugation at 4000 g for 20 minutes, formed a dark grey pellet. A volume of 1 L culture formed 12-19 g wet cell pellet. Expression of *Bc*R2b was adapted from Tomter et al. (Tomter et al., 2008) and the protein was expressed in a similar way as *Bc*NrdI. To express *Bc*R2b metal-free, ethylenediaminetetraacetic acid (EDTA) (PanReac AppliChem, Darmstadt, Germany) was added to the large-scale cultures shortly before induction with IPTG to a final concentration of 1 mM. A volume of 1 L culture gave about 8 g wet cell pellet. The pellets were flash-frozen in liquid nitrogen and stored at -20 °C until further use.

### Purification

#### *Bc*NrdI

The purification of NrdI was adapted from Røhr et al (Røhr et al., 2010). About 20 g of bacterial cell pellet was resuspended in lysis buffer (100 mM Tris-HCl pH 7.5) supplemented with a tablet of EDTA-free cOmplete Protease Inhibitor S2 Cocktail Tablet (PIC) (Roche, Solna, Sweden) and DNase (PanReac Applichem) was added. Cells were lysed with a sonicator, Sonics VCX130 (Sonics, Newtown, CT), and the soluble fraction separated from cell debris by centrifugation at 40000 g for 30 minutes at 4 °C. The NrdI protein was precipitated by slow addition of ammonium sulphate (NH_4_SO_4_) to the lysate to a final concentration of 60 % w/v (0.37 g/ml) while stirring at 4°C. The precipitate was pelleted by centrifugation at 20000 g for 20 minutes and 4°C and subsequently solubilised with a minimal volume of size exclusion chromatography (SEC) buffer (50 mM Tris pH 7.5). The protein was desalted by dialysis overnight at 4°C against dialysis buffer (10 mM Tris pH 7.5). The desalted protein was filtered through a 0.45 µm filter to remove precipitate and loaded onto a Q Sepharose High Performance 5 ml anion exchange column (Cytiva, Uppsala, Sweden), washed with SEC buffer and eluted with a gradient from 0-50% elution buffer (50 mM Tris-HCl pH 7.5, 1M KCl). The fractions containing the desired protein could be identified by the bright orange colour of NrdI (Fig. S1). Relevant fractions were pooled, concentrated with a Vivaspin 20 centrifugal concentrator with a 30,000 Da molecular weight cut-off polyethersulfone membrane (Vivaspin 30k concentrator) (Sartorius, Göttingen, Germany) and injected onto a HiLoad Superdex 75 prep grade size-exclusion column (Cytiva). Orange fractions were analysed for purity by sodium dodecyl sulphate–polyacrylamide gel electrophoresis (SDS-PAGE) and fractions with a 95% or higher purity were pooled. The theoretical molecular weight of NrdI was calculated to be 13449 Da with ProtParam (Gasteiger et al., 2005) and the extinction coefficient ε_447_ = 10.8 mM^-1^ cm^-1^ (Berggren et al., 2014) was used to determine the protein concentration using UV-vis spectroscopy. The protein was concentrated to 25 mg/ml, flash frozen in liquid nitrogen and stored at -80 °C until further use.

#### *Bc*R2b

The R2b purification protocol was adapted from Tomter et al. (Tomter et al., 2008). The protein was produced metal free; for that purpose, EDTA was included in the lysis buffer to inhibit metal uptake. About 20 g of bacterial cell pellet was resuspended in lysis buffer (100 mM Tris-HCl pH 7.5, 1 mM EDTA) with a tablet of PIC and lysed by sonication. Streptomycin sulphate was added to a total of 2.5% w/v and incubated for 10 minutes at 20° C for DNA precipitation. The crude lysate was cleared by centrifugation for 30 min at 40000 g. The supernatant was cleared from contaminants by adding 40 % (0.24 g/ml) NH_4_SO_4_ at 20°C followed by centrifugation at 20000 g for 20 minutes. The NH_4_SO_4_ concentration for the remaining supernatant was increased to 50 % (0.31 g/ml) at 20° to precipitate R2b. After the second centrifugation at 20000g for 20 minutes the pellet was dissolved with a minimal volume of lysis buffer. The protein was diluted 4x with buffer A (50 mM Tris pH 7.5, 1.5 M NH_4_SO_4_) to increase the salt concentration sufficiently. The diluted protein was filtered through a 0.45 µm membrane filter and loaded onto a column packed with 20 ml Phenyl Sepharose High Performance resin (Cytiva). The protein was washed with buffer A and eluted with a gradient with SEC buffer (50 mM Tris pH 7.5). Several column volumes were needed to elute the protein. The elution fractions were analysed by SDS-PAGE. Protein containing fractions were pooled, concentrated by a Vivaspin 50k concentrator and injected onto a HiLoad Superdex 200 prep grade column (Cytiva). Elution fractions were analysed by SDS-PAGE and pure fractions pooled. The theoretical molecular weight and the extinction coefficient of R2b were calculated with ProtParam to be 37017 Da and ε_280_ = 48.36 mM^-1^ cm^-1^ respectively (Gasteiger et al., 2005). R2b was concentrated to 50 mg/ml with a Vivaspin 50kDa concentrator, flash frozen and stored at -80 °C until further use. A total of 20 g bacterial cell pellet yielded about 100-150 mg pure protein.

### Quantification of R2b metal content with TXRF

To control the metal content of *Bc*R2b the protein was measured with total reflection x-ray fluorescence spectroscopy (TXRF) using a Bruker PicoFox S2 spectrometer (Bruker, Billerica, MA). Three independently prepared replicates of the concentrated protein at 1.7 mM were mixed 1:1 with a gallium standard at 20 mg/l and dried on top of a siliconized quartz sample carrier. The discs were individually measured before use to avoid external contamination of the sample. The results were analysed with the Bruker Spectra software version 7.8.2.0 provided with the instrument.

### Crystallization of the *Bc*R2b-NrdI complex

The *Bc*R2b-NrdI protein complex was prepared for crystallization as follows: 0.25 mM *Bc*R2b in 50 mM Tris-HCl pH 7.5 was incubated with 12 molar equivalents of MnCl_2_ for 10 minutes at 20° C, then 0.25 mM *Bc*NrdI was added and the mixture incubated for another 15 minutes at 20° C to ensure complex formation. New crystallization conditions were screened since the crystallization condition published in (Hammerstad et al., 2014) could not be reproduced. Initial hits in conditions C5 of the JSCG+ crystallisation screen (0.1 M Sodium Hepes pH 7.5, 0.8 M Sodium phosphate monobasic monohydrate,0.8 M Potassium phosphate monobasic) and the A7 condition of the PACT premier crystallization screen (0.1 M Sodium acetate pH 5.0, 0.2 M NaCl, 20 % w/v PEG 6000) (both Molecular Dimensions, Sheffield, UK) were further optimized to yield crystals between 10 µm and 50 µm in the longest axis (Fig. S1). The final crystallization protocol was established as described: Crystals spontaneously grew in a hanging drop vapour diffusion experiment after one to two days at 20°C. Crystallization condition A (0.1 M Hepes pH 7.0, 0.6-0.85 M sodium phosphate monobasic monohydrate, 0.6-0.85 M potassium phosphate monobasic) was manually mixed with the protein complex solution in a ratio of 1:1 (1 µl + 1 µl). The crystals grown in this experiment were used to produce microseeds for crystallization with the batch method. Two crystal containing drops were transferred into 50 µl of crystallization condition A in the bottom of a 1.5 ml microcentrifuge tube. A seed bead (Saint Gobain, Aachen, Germany) was placed into the solution, the crystal crushed by vigorous shaking and the mixture used as seed stock for the following crystallization experiments. Batch crystallization was set up in PCR tubes at 20 °C. A volume of 40 µl of crystallization condition B (0.1 M MnCl_2_, 0.1 M sodium acetate 5.0, 5% PEG 6000) were pipetted on top of 40 µl of protein complex solution. A volume of 8 µl of microseed solution was added to the tube and everything mixed by pipetting. Orange rhombohedron shaped crystals typically sized between 20 and 50 µm in the longest axis grew over night, forming an orange pellet at the bottom of the tubes. The crystals were resuspended and pooled in 2 ml microcentrifuge tubes (Fig. S1). Crystals used to collect the dataset for *Bc*R2b_MnMn_-NrdI_ox_ could be directly loaded into a Gastight SampleLock (Hamilton, Bonaduz, Switzerland) syringe for sample delivery. The crystals used for the *Bc*R2b_MnMn_-NrdI_hq_ dataset had to be chemically reduced first as described below.

### Reduction protocol

Crystals for the *Bc*R2b_MnMn_-NrdI_hq_ dataset were chemically reduced before data collection. All following steps were conducted in an anaerobic glove box with O_2_ below 10 ppm at room temperature. A volume of 900 µl of pooled crystal slurry was gently centrifuged, forming a dense crystal pellet. Of the supernatant 800 µl were carefully removed, collected separately and supplemented with 18 µl of freshly prepared, anaerobic 1 M sodium dithionite (DT). The DT-containing supernatant was gently mixed with the crystal pellet (final DT concentration of 20 mM) and the colour change of the crystals from a bright orange to a faint, light yellow was observed in a matter of minutes. The DT was subsequently washed out by gently spinning down the crystals, removing the supernatant without disturbing the crystal pellet, replacing it with anaerobic wash buffer (100 mM MnCl_2_, 50 mM sodium acetate pH 5.0, 2.5% (w/v) PEG 6000, 10% (v/v) glycerol) and gently resuspending the crystals. This washing step was repeated 3 times, adding wash buffer in the last step to a volume of 800 µl. The final concentration of DT in the crystal slurry was below 4 µM. All of the crystal slurry was loaded into a 1 ml gastight SampleLock Hamilton syringe and stored in the anaerobic box until sample injection.

### Data collection

The *Bc*R2b-NrdI crystals were initially tested for stability and diffraction quality at SACLA in Japan during experiment 2017B8085. The sample was delivered with a grease extruder setup installed on site (Sugahara et al., 2015; Tono et al., 2013). Hydroxyethyl cellulose matrix (Sugahara et al., 2017) and the protein crystals were mixed together in a volume ratio matrix to crystal pellet of 9:1 and the mixture ejected with a HPLC pump through a 150 µm nozzle with a flow rate of 1-1.5 µl/min, delivering the sample into the XFEL beam. X-rays were delivered as <10 fs long pulses at 10.9 keV with 30 Hz repetition rate and a typical pulse energy of around 0.32 mJ with a beam size of 2 × 2 µm^2^ (fwhm). The diffraction of the sample was recorded 100 mm downstream of the interaction point on an Octal MPCCD detector. The initial data collection showed stability of the crystals over days and a diffraction quality to about 2 Å. The datasets used in this paper were collected at LCLS, California during experiment LU50. X-ray pulses at 9.5 keV with a pulse energy of 4 mJ, 30 Hz repetition rate and a duration of around 35 fs were generated and used for X-ray diffraction in the macromolecular femtosecond crystallography (MFX) experimental hutch (Sierra et al., 2019); the diffraction was recorded on a Rayonix MX340 (Rayonix L.L.C, Evanston, USA) detector. The sample was delivered into the X-ray interaction point with the drop-on-tape method; the detailed method was described by Fuller et. al (Fuller et al., 2017). Briefly, crystal slurry in a 1 ml SampleLock syringe was pumped with a syringe pump (KD scientific, Holliston, MA) at a flow rate of 8-9 µl/min through a silica capillary into a 6 µl sample reservoir. An acoustic transducer transferred crystal containing droplets of 2.5-4 nl volume onto a Kapton tape, which transported the droplets into the X-ray beam at 28° C and 27% relative humidity with a speed of 300 mm/s, which resulted in the crystals being exposed to the He-environment for about 0.8 s. The enclosure of the setup was filled with a He-atmosphere with an O_2_ level below 0.1 % for the *Bc*R2b_MnMn_-NrdI_hq_ data collection.

### Processing, Structure Determination and Refinement

The datasets for *Bc*R2b_MnMn_-NrdI_hq_ and *Bc*R2b_MnMn_-NrdI_ox_ were processed with cctbx.xfel (Brewster, Young, et al., 2019; Hattne et al., 2014; Sauter, 2015) and DIALS (Brewster et al., 2018; Winter et al., 2018). We performed joint refinement of the crystal models against the detector position for each batch to account for small time-dependent variations in detector position and also corrected for the Kapton tape shadow (Fuller et al., 2017). Data was scaled and merged to 2.0 Å resolution using cctbx.xfel.merge with errors determined by the ev11 method (Brewster, Bhowmick, et al., 2019) based on CC_1/2_ and multiplicity values and on R-factors after initial refinement. Data statistics are available in Table 1. Both structures were solved by molecular replacement and refined independently with the PHENIX Suite (Liebschner et al., 2019). The phases were solved with phenix.phaser (McCoy et al., 2007). A *Bc*R2b-NrdI complex structure (PDB ID: 4BMO (Hammerstad et al., 2014)) was modified by manually removing waters and alternate conformations and used as a starting model. The suggested solutions were in the same space group (*C*222_1_) as the starting model with similar unit cell dimensions (Table 1) with one complex in the asymmetric unit. The *R*_free_ set of the 4BMO model corresponding to 5% of reflections was assigned to all datasets with phenix.reflection_tools. Restraints of the entry “FMN” of the ligand database provided by phenix were used for the oxidized FMN in *Bc*R2b_MnMn_-NrdI_ox_. Restraints for the hydroquinone FMN in *Bc*R2b_MnMn_-NrdI_hq_ were generated with phenix eLBOW (Moriarty et al., 2009). The datasets were iteratively refined with phenix.refine (Afonine et al., 2012), examined and built in coot (Emsley et al., 2010) and validated with MolProbity (Williams et al., 2018). Refinement of all atoms included isotropic B-factors, TLS parameters, occupancy and reciprocal space refinement with a high-resolution cut off of 2.0 Å. Waters were initially added using phenix.refine and in later refinements corrected manually. The metal occupancy for the reduced structure was fixed manually after several cycles of refinement. An overview over refinement and model quality statistics can be found in Table 1, created with phenix.table_one. The refined structures were compared and RMSD values calculated with SSM superimpose (Krissinel & Henrick, 2004).

### Figures

Molecular figures were prepared with the PyMOL Molecular Graphics System, Version 2.4.2 Schrödinger, LLC. Scheme 2 was designed with ChemDraw (PerkinElmer Informatics). The bar graph in Fig. 3 A was generated with GraphPad Prism, Version 9.2.0 for iOS (GraphPad Software, San Diego, California USA). All figures and schemes except Scheme 2 were designed and assembled in Affinity Designer (Serif (Europe) Ltd, Nottingham, UK)

## Acknowledgements

We thank Robert Bolotovsky, Iris D. Young and Lee J. O’Riordan for help with data processing and the staff at LCLS and SACLA. We thank Kristine Grāve for input and helpful discussions during data analysis and manuscript writing.

This work was supported by the Swedish Research Council (2017-04018 and 2021-03992 to M.H.), the European Research Council (HIGH-GEAR 724394 to M.H.), the Knut and Alice Wallenberg Foundation (20217.0275 and 2019.0436 to M.H.), and National Institutes of Health grants GM133081 (to K.D.S.), GM117126 (to N.K.S.), GM55302 (to V.K.Y.), GM110501 (to J.Y.) and GM126289 (to J.K.). A.M.O., P.A., and A.Bu. were supported by Diamond Light Source, the UK Science and Technology Facilities Council (STFC), a jointly funded strategic award from the Wellcome Trust and the Biotechnology and Biological Sciences Research Council (102593 to James Naismith), a Wellcome Investigator Award (210734/Z/18/Z to A.M.O.), and a Royal Society Wolfson Fellowship (RSWF\R2\182017 to A.M.O.). The DOT instrument used in this research was funded by Department of Energy (DOE), Office of Science, Office of Basic Energy Sciences (BES), Division of Chemical Sciences, Geosciences, and Biosciences (to J.K., J.Y., and V.K.Y.). XFEL data was collected under proposal LU50 at LCLS, SLAC, Stanford, USA, and under proposal 2017B8085 at BL2 of SACLA, Japan. The Rayonix detector used at LCLS was supported by the NIH grant S10 OD023453. Use of the LCLS, SLAC National Accelerator Laboratory, is supported by the U.S. DOE, Office of Science, BES, under contract no. DE-AC02-76SF00515.

## Author Contributions

J.J., H.L., V.S., O.A. and M.H. developed the study

C.P., I-S.K., S.G., P.S.S., A.M.O., F.D.F., and J.K. developed, tested and ran the sample delivery system

A.S.B., M.D., K.D.S., A.Bh., A.Bu., and N.K.S. processed and analysed XFEL data F.D.F., A.Ba. operated the MFX instrument

J.J., H.L., O.A., M.H., C.P., I-S.K., A.S.B., S.G., K.D.S., A.Bh., P.S.S., A.Bu., P.A., A.M.O., F.D.F.,

A.Ba., N.K.S., V.K.Y., J.Y., J.K. performed the LCLS experiment

V.S., H.L., P.A., F.D.F., M.H., M.H.C., A.M.O., S.O., K.T., V.K.Y., J.K., J.Y. performed the SACLA experiment

J.J. and H.L. produced and purified proteins and developed the crystallization protocol

J.J. refined, analysed and interpreted the data and wrote the manuscript with input from all authors

H.L. and M.H. supervised, edited and revised the manuscript

## Data availability

The atomic coordinates and crystallographic data have been deposited in the Protein Data Bank (https://www.pdb.org/) under the following accession codes: *Bc*R2b_MnMn_-NrdI_ox_: 7Z3D; *Bc*R2b_MnMn_-NrdI_hq_: 7Z3E.

## Supplementary Information

**S 1.**
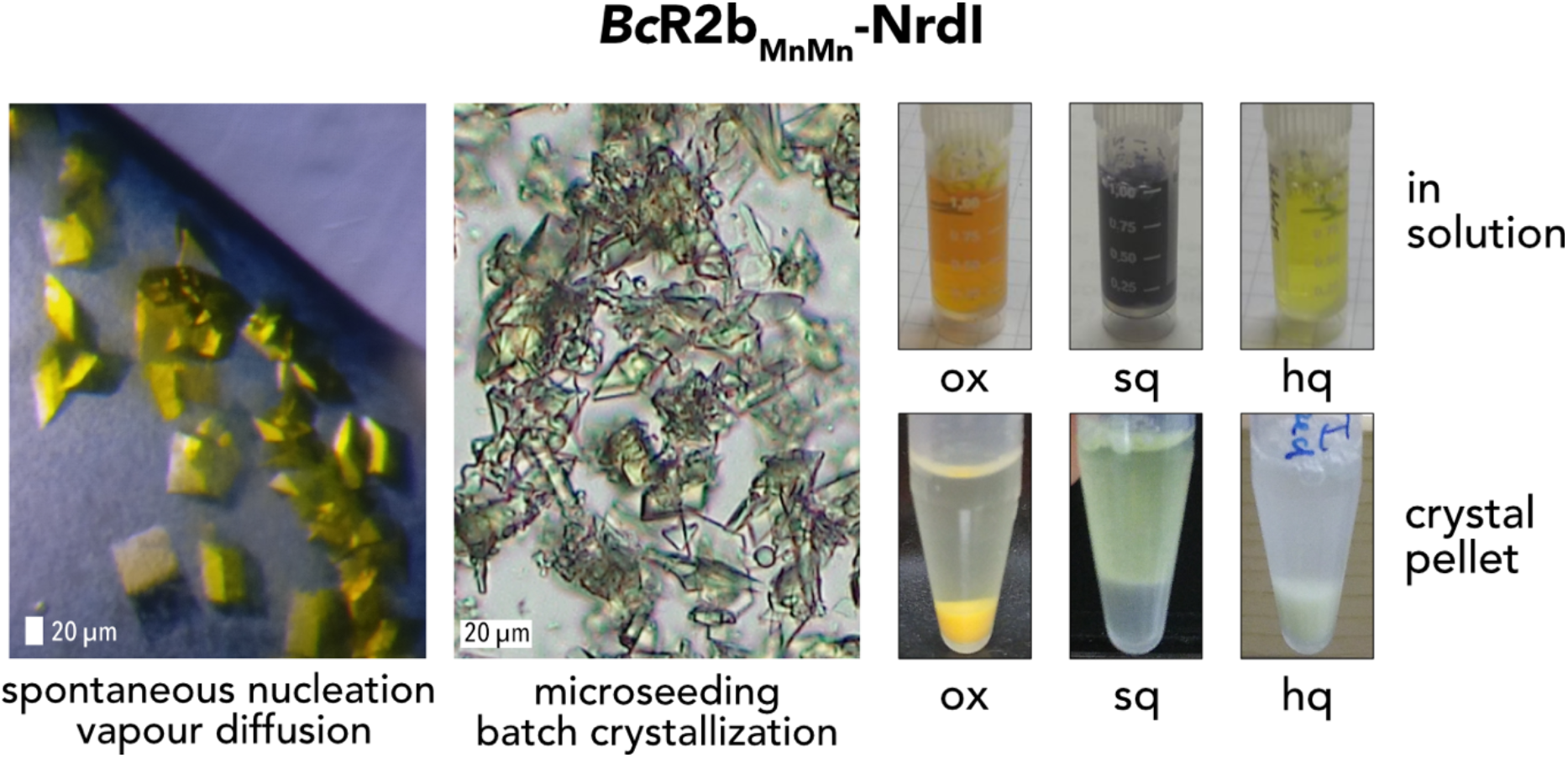
Crystallization of the di-manganese *Bc*R2b-NrdI complex. Spontaneous nucleation yields crystals of the length of 100 μm or longer. In the batch crystallization, crystals are significantly smaller with most crystals between 20 and 50 μm. For the two pictures on the left side, crystals of the complex with NrdI in the oxidized state are viewed under an optical microscope. On the right side, pictures are shown for the complex in solution (top row) and in crystals (bottom row) with NrdI in the oxidized, semiquinone and hydroquinone states. The oxidation state of FMN changes the colour of the complex in both solution and crystals.

**S 2.**
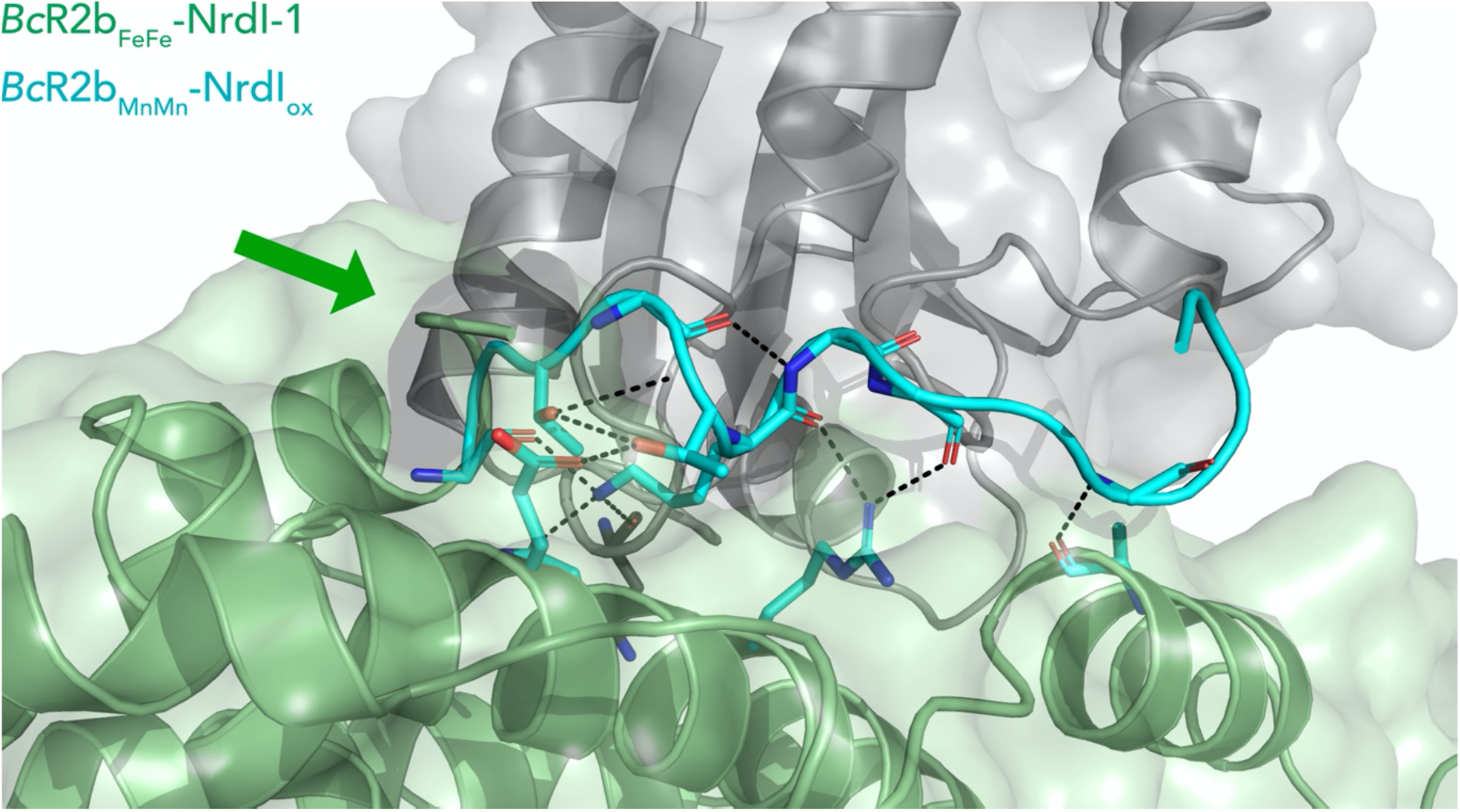
Extended C-terminus in *Bc*R2b_MnMn_-NrdI_ox_. Superposition of *Bc*R2b_FeFe_-NrdI-1 (PDB ID: 4BMO) (pale green for R2b and grey for NrdI) and *Bc*R2b_MnMn_-NrdI_ox_ (in cyan) showing an extension of the ordered C-terminus by 9 amino acids. The green arrow points towards the C-terminus of *Bc*R2b_FeFe_-NrdI-1. The C-terminus is situated in a groove between R2b and NrdI; hydrogen bonds are shown with black dashed lines and bonded side chains are shown as sticks.

**S 3.**
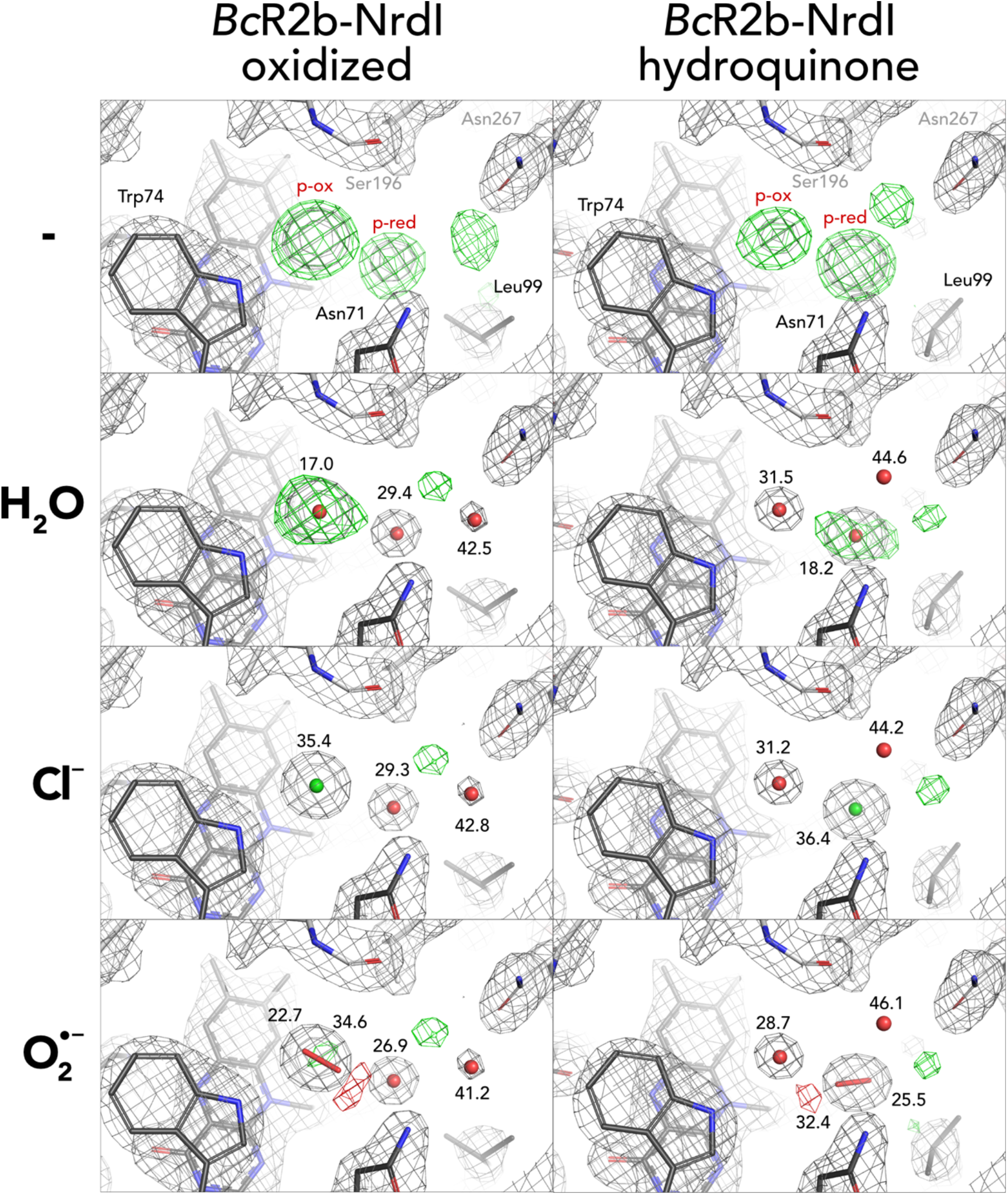
Different modelling results for unidentified density close to FMN. Results for both oxidized and hydroquinone *Bc*R2b-NrdI complex structures. 2Fo-Fc maps contoured at 2 σ are shown in grey, and Fo-Fc maps contoured at 4 σ with positive and negative density are shown in green and red, respectively. Refined B-factor values in Å^2^ are indicated for the different atoms.

**S 4.**
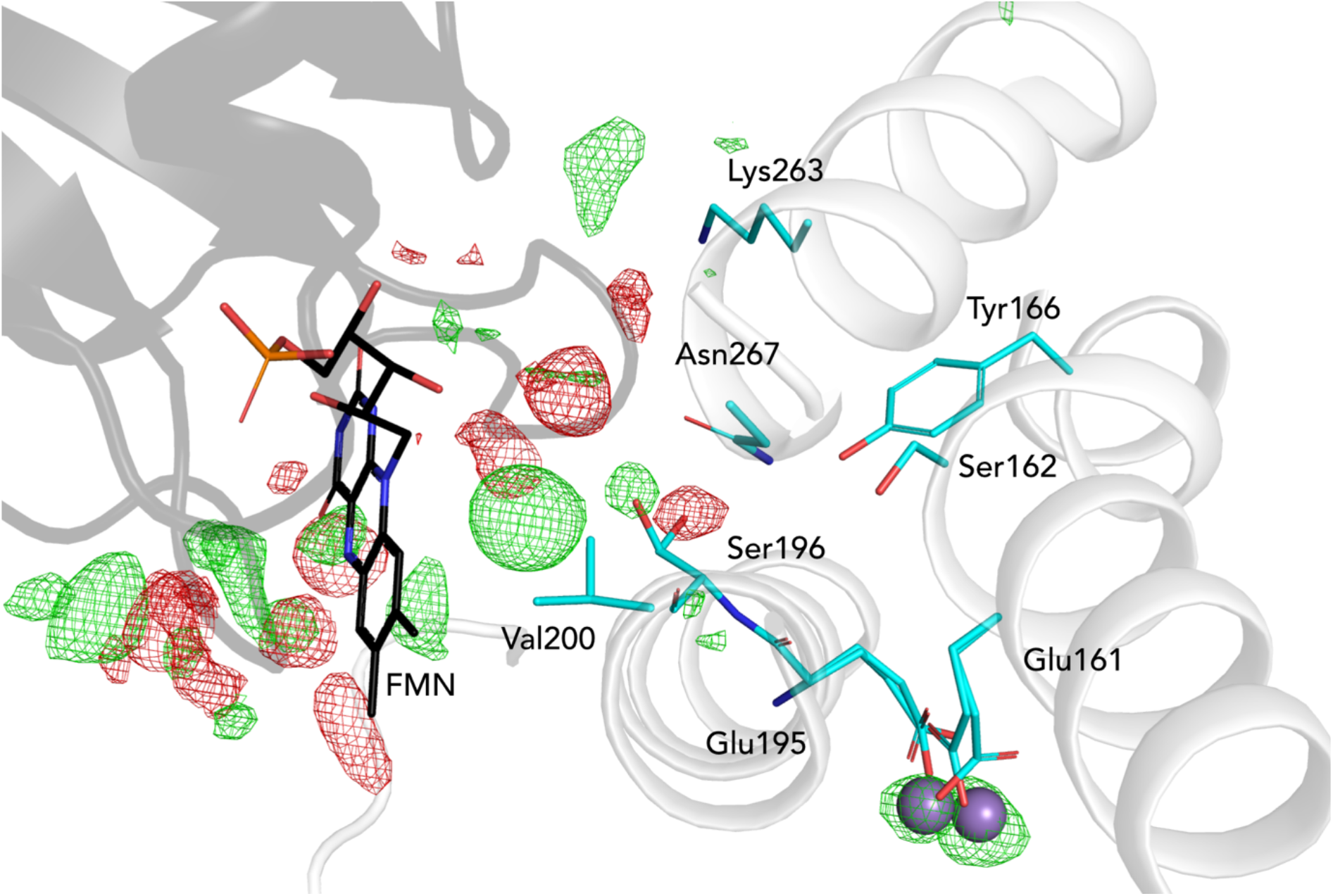
Channel between R2b and NrdI. NrdI is represented in black and R2b in white. Residues lining the channel from FMN (black sticks) towards the di-manganese site (purple spheres) are shown as sticks in cyan. The Fo(ox)-Fo(hq) map is shown as mesh and contoured at 4.5 σ with green as positive and red as negative density. The observed unknow molecule is in the vicinity of the channel. Changes around the FMN do not protrude towards the metal site in R2b.

**S 5.**
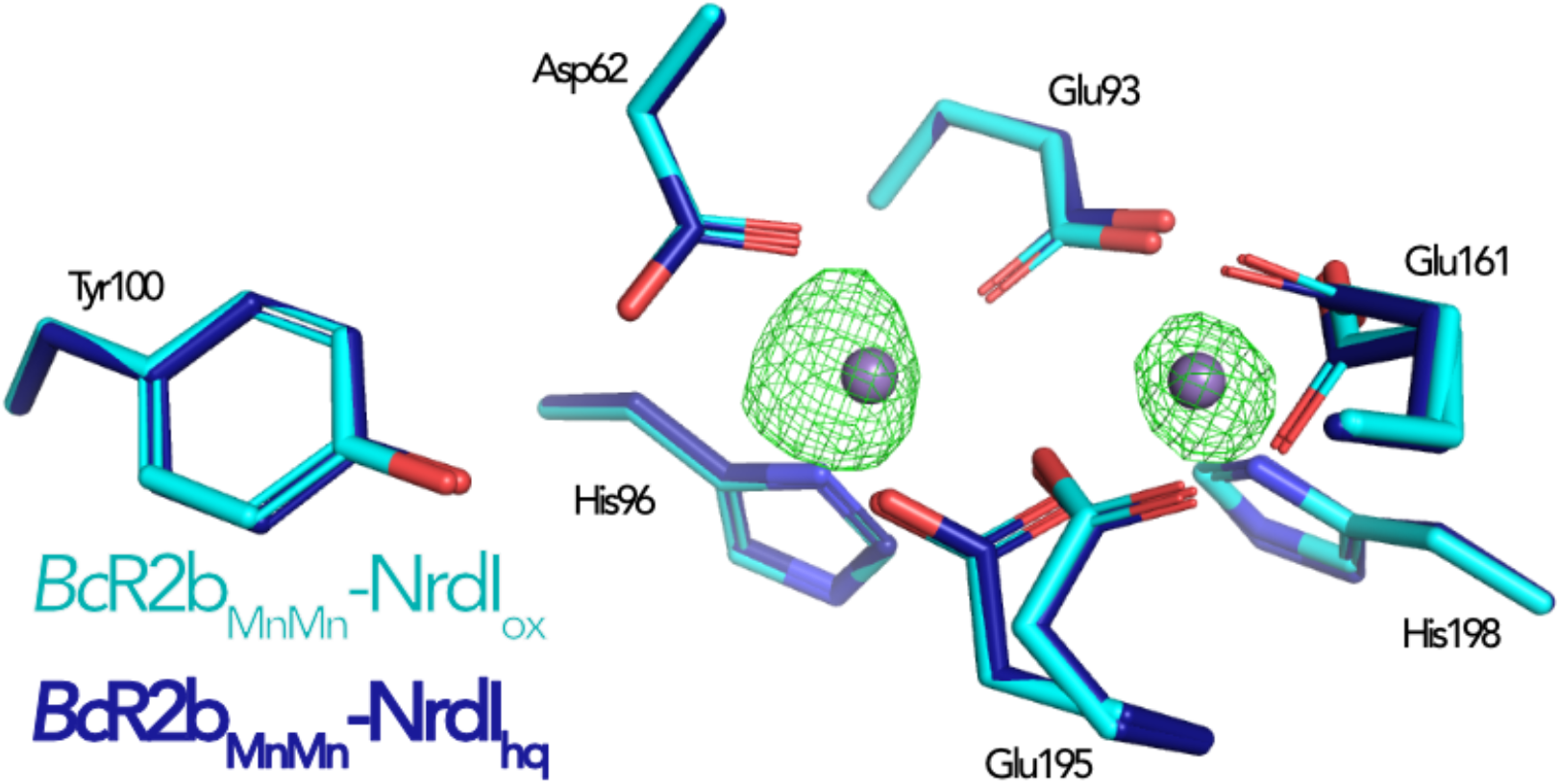
Comparison of the active site in di-manganese *Bc*R2b_MnMn_-NrdI structures. The Fo(ox)-Fo(hq) map is shown as mesh and contoured at 4.5 σ. The only observable change between the structures is the metal occupancy.

